# How ecosystems recover from pulse perturbations: A theory of short- to long-term responses

**DOI:** 10.1101/115048

**Authors:** J.-F. Arnoldi, A. Bideault, M. Loreau, B. Haegeman

## Abstract

Quantifying stability properties of ecosystems is an important problem in ecology. A common approach is based on the recovery from pulse perturbations, and posits that the faster ecosystems return to their pre-perturbation state, the more stable they are. In theoretical studies the recovery dynamics are often collapsed into a single quantity: the long-term rate of return, called asymptotic resilience. However, empirical studies typically measure the recovery dynamics at much shorter time scales. In this paper we explain why asymptotic resilience is rarely representative of the short-term recovery. First, we show that, in contrast to asymptotic resilience, short-term return rates depend on features of the perturbation, in particular on the way its intensity is distributed over species. We argue that empirically relevant predictions can be obtained by considering the median response over a set of perturbations, for which we provide explicit formulas. Next, we show that the recovery dynamics are controlled through time by different species: abundant species tend to govern the short-term recovery, while rare species often dominate the long-term recovery. This shift from abundant to rare species typically causes short-term return rates to be unrelated to asymptotic resilience. Finally, we discuss how these findings might help to better connect empirical observations and theoretical predictions.

## Introduction

Ecosystem stability, or the way in which an ecosystem responds to a perturbation, is a long-standing topic of interest in ecology (May, 1973; Pimm, 1984; Tilman & Downing, 1994). Ecologists have used a variety of procedures to quantify ecosystem stability, differing in the characteristics of the perturbation and in the way the system response is measured. The perturbation can consist of a change in an environmental parameter (e.g., temperature, nutrient supply), which can last for a short or a long time. Alternatively, the perturbation can correspond to biomass addition or removal, which can be applied once or repeatedly. The system response can be assessed immediately after the perturbation or after a longer period of time, by measuring the overall state of the ecosystem or an ecosystem variable of specific interest (e.g., total biomass, nutrient uptake). This multitude of procedures has led to an overabundance of stability measures, whose relationships are often unclear (Grimm & Wissel, 1997; Ives & Carpenter, 2007; Donohue *et al.,* 2013).

In this paper we focus on one family of stability measures, namely those based on the ecosystem response to a pulse perturbation (i.e., a perturbation of relatively short duration; Bender *et al.,* 1984). We assume that the perturbation is not too strong so that after a sufficiently long time the ecosystem settles in a state that is comparable to the pre-perturbation state. The way the system returns to this state, which we call equilibrium, is then used to quantify the ecosystem's stability. Indeed, it is intuitive to assume that an ecosystem is more stable if it returns more quickly to its equilibrium. Several stability measures can be defined, differing in the time at which, and the ecosystem variable of which, the return to equilibrium is assessed. Terms used for these stability measures include return time, recovery rate, and resilience. The term resilience, however, might lead to confusion, because it is also used for a rather different set of stability measures (Holling, 1973; Gunderson, 2000).

The approach to quantify ecosystem stability based on the return to equilibrium is popular in both empirical and theoretical studies. Pulse perturbations are an appropriate model for many natural disturbances, such as floods, forest fires and disease outbreaks. They have also been widely applied in experimental ecosystems. These studies typically concentrate on the short-term return to equilibrium, due to practical difficulties of collecting time series data over a long time period (e.g., Steiner *et al.,* 2006; Downing & Leibold, 2010; Hoover *et al.,* 2014; Wright *et al.,* 2015). This stands in sharp contrast with theoretical work, in which the return to equilibrium is mainly studied at long time scales (e.g., Rooney *et al.,* 2006; Loeuille, 2010; Thébault & Fontaine, 2010; Tang *et al.,* 2014). This is due to the fact that the longterm rate of return to equilibrium can be easily computed. In particular, this rate, known as asymptotic resilience, can be expressed in terms of an eigenvalue of the community matrix (we recapitulate this theory in the next section). Hence, while the return to equilibrium is clearly of common interest, theoretical and empirical studies focus on different aspects.

Here we investigate the discrepancy between (empirical) short-term and (theoretical) longterm return to equilibrium. The problem that ecological theory and data do not necessarily address the same time scales has been emphasized before (reviewed in Hastings, 2010). Of particular relevance is the work of Neubert & Caswell (1997), who argued that the initial response of an ecosystem to a pulse perturbation can strongly differ from its long-term response. They described ecosystems that eventually return to equilibrium for any perturbation (i.e., stable ecosystems) but initially move away from equilibrium following some perturbations (called reactive systems; see below for more details). Our work can be seen as an extension of Neubert & Caswell’s theory. Specifically, while their work dealt with the perturbation that causes the strongest response, we study how the ecosystem recovers on average after a perturbation. As we argue, the latter approach is more relevant from an empirical viewpoint. We come back to the relationship between Neubert & Caswell’s and our paper in the discussion section.

We start by proposing a precise definition of return rates and return times, covering the range from very short term (initial response) to very long term (asymptotic response). Then, we show that short-term and long-term return rates differ in their dependence on the perturbation direction, i.e., the way its intensity is distributed over species. This dependence can be strong for short times, but vanishes in the limit of very long times (i.e., asymptotic resilience). We propose to summarize the distribution of return rates for different perturbations by its median, for which we present a simple and accurate approximation. This allows us to compare shortterm and long-term return rates on an equal footing. Then, using these return rates, we find that species abundance can play a predominant role in the recovery dynamics. In particular, rare species often have a strong effect on the long-term response, while their effect on the shortterm response is mostly limited. We describe the mechanism behind this observation, and illustrate its genericity using a random many-species community model. The model results show that asymptotic resilience and short-term return rates are typically disconnected. This suggests that asymptotic resilience provides only a partial view on the recovery dynamics, and that empirically more relevant predictions might be obtained from short-term return rates, such as those introduced and studied in this paper.

## Defining return rates and return times

The terms resilience, return rate and return time are widely used to describe the recovery of an ecosystem after a pulse perturbation. Especially in empirical studies, these stability measures are often constructed in an *ad hoc* manner. This has led to a variety of mostly incompatible definitions. Here we propose a framework in which at least some of these measures can be incorporated.

The study of the recovery dynamics starts by specifying a reference state of the ecosystem. This is the state from which the ecosystem is perturbed and to which it returns after the perturbation. Typically, the reference state corresponds to a dynamic equilibrium, characterized by relatively small fluctuations around a fixed average. The pulse perturbation then induces a much larger displacement, such that the ecosystem leaves its reference state. This initiates the recovery dynamics, which last until the ecosystem reaches the reference state again. At that time the displacement induced by the perturbation has been absorbed by the ecosystem.

It is practically impossible to study the recovery once the displacement induced by the pulse perturbation has become indistinguishable from the fluctuations that characterize the dynamic equilibrium. However, almost all previous theoretical work has focused on the long-term return to equilibrium, which is observable only if the equilibrium fluctuations are absent (or very small). It is customary to model the reference state as a static equilibrium without fluctuations (May, 1973, 1974). We also adopt this approach, but we emphasize that our results on the short-term recovery also hold for a fluctuating reference state.

In short, we model the reference state as an equilibrium of a deterministic dynamical system. We assume that this equilibrium is stable. Here the term ‘stable’ is used in the sense of the stability criterion, that is, the ecosystem returns to equilibrium for any sufficiently small displacement from equilibrium. Note that this does not exclude the possibility that larger displacements can push the ecosystem to a different state (another equilibrium or a more complex attractor).

Moreover, we restrict attention to small displacements from equilibrium, which corresponds to weak pulse perturbations. This assumption allows us to rely on the theory of linear dynamical systems, which has provided a benchmark in the study of ecological stability since May's (1973) seminal work. Denoting the vector of dynamical variables (e.g., the biomass of the species in the ecosystem) by *N*(*t*) and the equilibrium by *N*\*, we construct the dynamics for the displacement vector *x*(*t*) = *N*(*t*) – *N*\*. These linearized dynamics are governed by the community matrix A, that is, the Jacobian of the non-linear dynamical equations evaluated at the equilibrium,

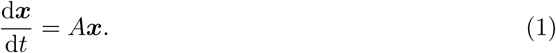

Equipped with the near-equilibrium dynamics of the unperturbed ecosystem, we can describe its response to a pulse perturbation. We assume that the perturbation is applied at time *t* = 0 and that the system was previously at equilibrium (i.e., *x*(*t*) = 0 for *t* < 0). The pulse perturbation is characterized by a vector *u* and describes the ecosystem’s state immediately after the perturbation (i.e., *x*(0^+^) = *u*). Using equation (1) we find that the displacement *x*(*t*) changes as,

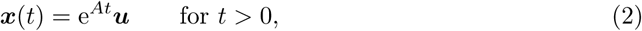

where e^A^ denotes the matrix exponential of *A*. Because we assumed the equilibrium to be stable (in the sense of the stability criterion), the system returns to equilibrium after a sufficiently long time *t*, that is, lim_t→∞_ *x(t)* = 0.

We are interested in quantifying how stable the system is, based on the idea that a more stable system returns faster to equilibrium. This general idea can be implemented in several ways, leading to different stability measures, as we will show below. Here we introduce one stability measure that will serve as a reference throughout the paper. It is based on the asymptotic return to equilibrium,

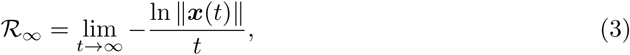

where the Euclidean norm 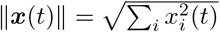 measures the distance to equilibrium. Equation (3) states that ||x(t)|| decays asymptotically as e^−𝓡_∞_*t*^. Note that, in principle, 𝓡_∞_ depends on the perturbation vector *u* = *x*(0^+^). However, it can be shown that 𝓡_∞_ is the same for virtually any perturbation *u* (see Appendix A). This common value, called asymptotic resilience, is equal to −ℛℯ(λ_dom_(*A*)), where λ_dom_(*A*) is the dominant eigenvalue of the community matrix *A*, i.e., its eigenvalue with the largest real part (ℛℯ(λ) denotes the real part of the complex number λ). Recall that ℛℯ(λ_dom_(A)) < 0 for stable systems (in the sense of the stability criterion), so that 𝓡_∞_ = −ℛℯ(λ_dom_(A)) is positive.

### Return rates

While the asymptotic return to equilibrium leads to a stability measure with elegant mathematical properties, only the return to equilibrium at finite times is of practical interest. We define two finite-time return rates: the instantaneous return rate at time *t*,

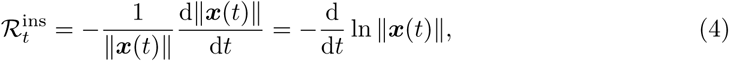
and the average return rate in the interval [0, t],

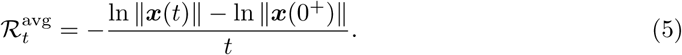

To illustrate this definition, we apply it to a system consisting of a single species. In this case the community matrix is a number, which has to be negative for the system to be stable (in the sense of the stability criterion). Hence, for *A =* −α with *α* > 0, we find *x(t) = u*e^−*α*t^ and 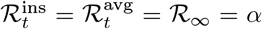 for all times *t*. Stability measures 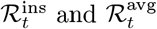 are expressed in units of reciprocal time, like asymptotic resilience.

The construction of return rates 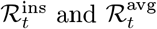 is illustrated in Fig. 1. We apply a pulse perturbation to a two-species community at equilibrium. From the recovery dynamics of the biomass variables *N*_1_*(t)* and *N*_2_*(t),* we obtain the distance to equilibrium ||x(t)|| as a function of time (panel A). To construct the return rates 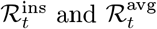, we represent this distance to equilibrium on a logarithmic scale (panel B). The instantaneous return rate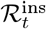 at time *t* is obtained as (minus) the slope of this curve at time *t*. The average return rate 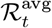 at time t is obtained as (minus) the slope of the line connecting the distances to equilibrium at times 0 and t. Note that the two return rates can differ strongly; they can even have opposite sign. For example, at time *t* ≈ 1.2, we have 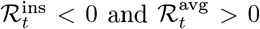. The former inequality means that the trajectory moves away from equilbrium at that time, while the latter inequality means that since the end of the perturbation the trajectory has come closer to equilibrium.

**Figure 1:**
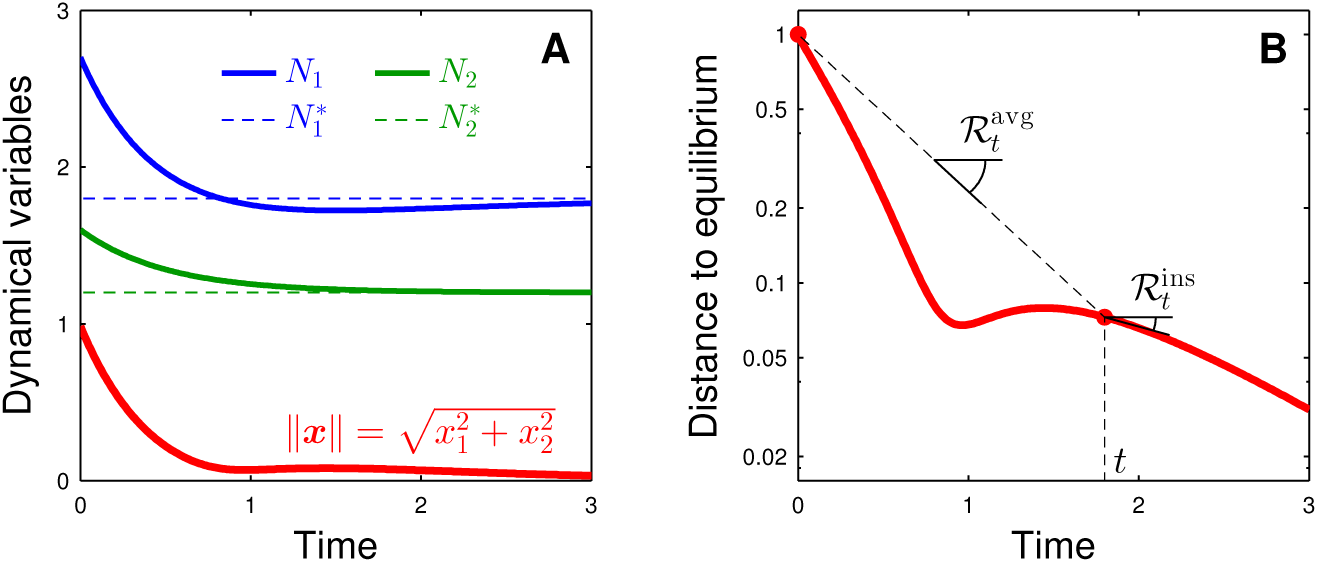
Definition of return rates. The response of an ecological system to a pulse perturbation contains information about the system’s stability, as illustrated here for a system of two interacting species. Panel A: We apply a pulse perturbation after which the species biomass *N*_1_(*t*) (blue) and *N*_2_(*t*) (green) return to their equilibrium values 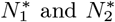. We monitor the recovery dynamics by the distance to equilibrium (red), 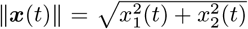 with x_i_(t) = N_i_(t) − N_i_^*^. Panel B: The relative rate at which the distance to equilibrium diminishes is a commonly used stability measure (note the logarithmic scale on the *y*-axis). Here we distinguish between the average rate of return 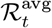 over the period [0, *t*], and the instantaneous rate of return 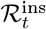 at time t. These two measures can largely differ, and can even have opposite sign. Parameter values: *N*^*⊤^ = (1.8,1.2), 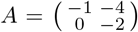 and *u*^⊤^ =(0.9, 0.4).

It is instructive to compare the behavior of return rates 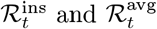 for very small and very large times *t*. It holds generally that 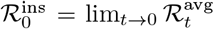, which follows directly from definitions (4) and (5). These definitions can also be used to show that 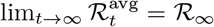. However, the analogous relationship 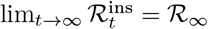 does not hold generally. It is satisfied for the example of Fig. 1, but it is not for the one of Fig. A.1. In the latter, return rate 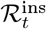 continues to oscillate between positive and negative values for large time *t*, so that 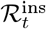 does not tend to a steady value for large time *t* (technically, 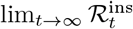 does not exist). This problem can be avoided by using a time-averaged version of 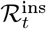, such as 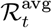. This is one reason why we focus on the average return rates 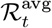 in this paper.

Definitions (3), (4) and (5) use the distance to equilibrium ||x(t)|| to quantify the extent of the displacement. Other stability measures can be constructed by using other scalar quantities derived from the displacement vector *x(t).* For example, by considering the sum Σ*_i_x_i_(t)*, one can study the return dynamics of total biomass, as has been used to address ecosystem-level stability (Tilman, 1996). While the theory in this paper is based on the distance to equilibrium, it can be extended to other ecosystem variables (see Appendix B).

### Return times

Return rates measure the speed at which an ecosystem approaches equilibrium. In practice, it might be more interesting to consider return times, that is, the time it takes for an ecosystem to recover from a perturbation. It is intuitively clear that return rates and return times are related. In this section we describe this link and point out some caveats.

Return time is defined as the amount of time between the perturbation and the instant at which the distance to equilibrium becomes smaller than a prespecified bound. To obtain an unambiguous definition, we also require that the distance to equilibrium remains smaller than that bound for all later times. In Appendix C we show that this definition leads to a set of return times parameterized by the bound on the distance to equilibrium, and we describe how these return times are related to the set of return rates 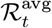. This provides another reason why in this paper we put the main focus on the average return rates 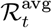. Note that, if the bound is chosen as the typical extent of the fluctuations in the equilibrium state, then the return time corresponds to the time during which the ecosystem response to the perturbation can be distinguished from the equilibrium fluctuations.

In theoretical studies the return time is often approximated as the reciprocal of asymptotic resilience. This approach, initiated by Pimm & Lawton (1977, 1978), is not self-evident, because it uses the asymptotic regime to describe the entire recovery dynamics, including the part shortly after the perturbation. That is, the approach implicitly assumes that the asymptotic return rate is a good proxy for the return rates at shorter times. As we argue extensively below, this need not be the case. This suggests that it is more appropriate to quantify the return time as the reciprocal of a finite-time return rate (recall that return rates have units of one over time). Again, the average return rate 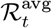 is particularly well suited, because it is based on the same part of the recovery dynamics as the return time, i.e., the period from the perturbation to some later, finite time.

In short, return times and return rates are related stability notions because they are derived from the same recovery trajectory. We have emphasized that different parts of the trajectory are used to approximate return times. Therefore, our study of the relationship between shortterm and long-term return rates is also directly relevant to the quantification of return times.

## Return rates depend on perturbation direction

There is a fundamental difference between asymptotic resilience and finite-time return rates. As mentioned above, virtually any pulse perturbation leads to the same asymptotic rate of return to equilibrium. Due to this remarkable property, asymptotic resilience has been called an intrinsic stability measure (Arnoldi *et al.,* 2016). In contrast, finite-time return rates do depend on features of the perturbation; they are not fully determined by the system dynamics alone. In this section we investigate this qualitative difference.

To do so, we again consider linear dynamical systems, characterized by the community matrix A. We apply a pulse perturbation, characterized by a perturbation vector u, or equivalently, by an initial displacement *x*(0^+^). Due to linearity, the perturbation intensity, quantified by the norm ||u||, has a trivial effect: when the perturbation is multiplied by a constant factor *α (u* ⟶ *α u*), the response *x(t)* is also multiplied by that factor *(x(t)* ⟶ *α x(t))*. This constant factor does not affect stability measures such as 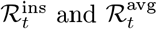 and (but it does affect return times). Hence, we only have to study the dependence of the recovery dynamics on perturbations vectors u with ||*u*|| = 1, i.e., the dependence on the perturbation direction. In ecological terms, the direction *u* defines the way the perturbation is distributed over the constituent species of the ecosystem.

Here we focus on the average return rate 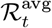 (recall that the term ‘average’ refers to the time average over the interval from 0 to t). The results are similar for the other stability measures introduced in the previous section. Recall that 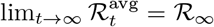, which is equal to asymptotic resilience. We denote the initial return rate by 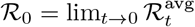.

To introduce our analysis, we look at a simple example of two non-interacting species (Fig. 2). The community matrix 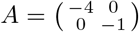 indicates that the first species responds four times faster to a displacement than the second species. The species with the slowest recovery determines asymptotic resilience R_∞_ = 1. This implies that for an arbitrary perturbation the system eventually returns to equilibrium with unit rate (Fig. 2B). However, this asymptotic rate of return is not informative about the short-term recovery. In particular, the system absorbs a perturbation that mainly affects the first species (perturbation ‘a’ in Fig. 2) much faster than a perturbation that mainly affects the second species (perturbation ‘b’ in Fig. 2).

**Figure 2:**
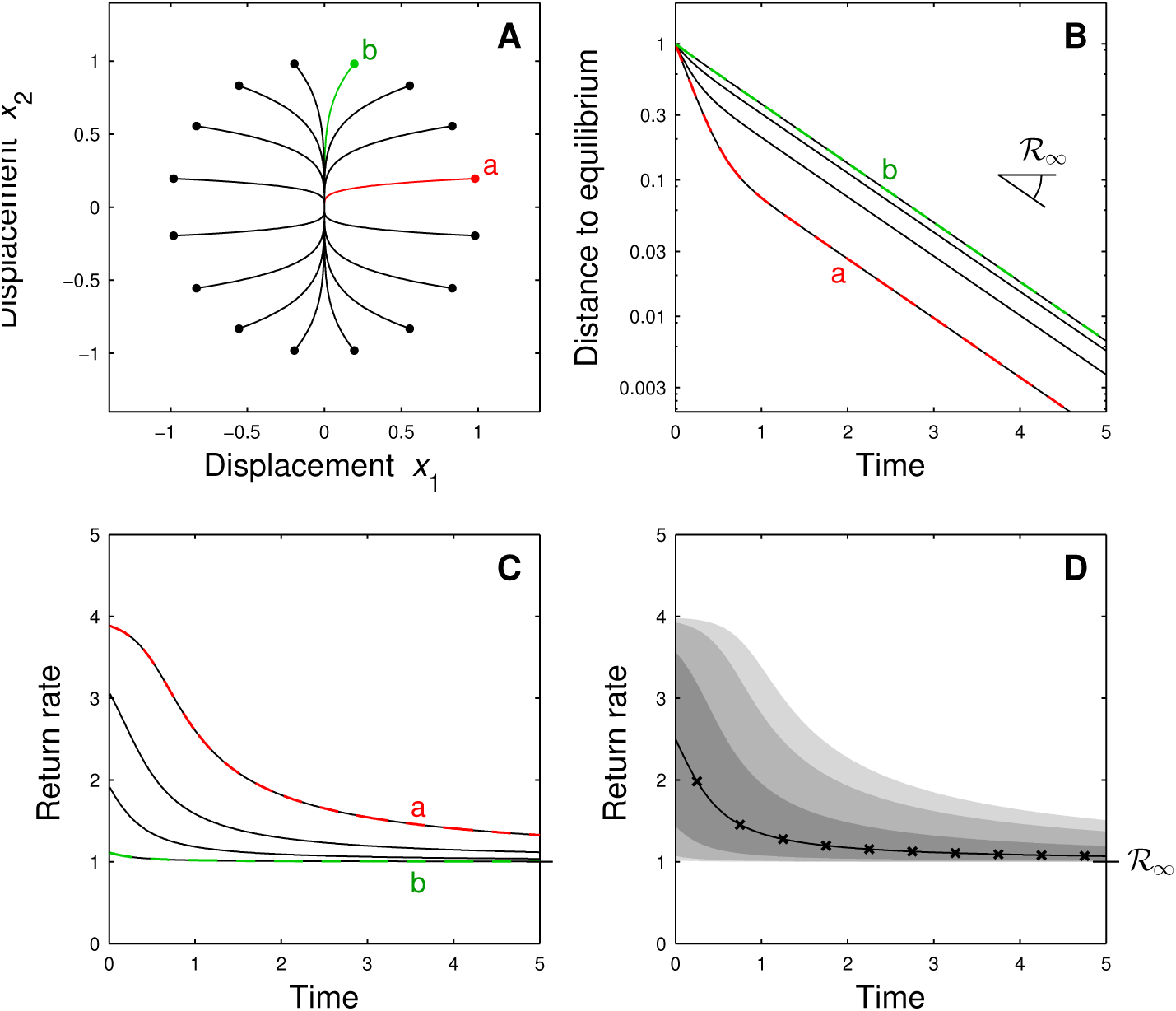
Return to equilibrium depends on perturbation direction – non-reactive case. Two-species system with community matrix 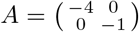, that is, species 1 responds four times faster than species 2. Panel A: Phase-plane trajectories for several perturbations *u* ***=*** *x*(0^+^). Perturbation ‘a’ (red) affects mostly species 1, while perturbation ‘b’ (green) affects mostly species 2. Note that all perturbations have the same intensity ||u|| = 1. Panel B: Dynamics of distance to equilibrium ||*x*(*t*)||. The return to equilibrium is faster for perturbation ‘a’ than for perturbation ‘b’. For all perturbations the distance to equilibrium eventually decays at a rate given by asymptotic resilience R_∞_. Panel C: Return rate 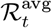 as a function of time *t*. As expected, the return rates are initially almost four times larger for perturbation ‘a’ than for perturbation ‘b’. Panel D: Statistics of return rate 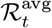 for random perturbations (fixed intensity, uniformly distributed). Full line: median computed from simulations; ×-marks: analytical approximation for median; shades of gray: 5%, 10%, 25%, 75%, 90% and 95% percentiles.

As a result, we find that return rates 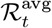 have a broad distribution (for different perturbations) at short times *t*, which becomes increasingly narrow at longer times *t* (see Fig. 2D). Asymptotic resilience, which is the lower limit of each of these distributions, is not a good predictor of the short-term return rate for an arbitrary perturbation. Better predictions can be obtained by looking at, e.g., the average of the return rate distribution. For a technical reason explained in Appendix D, we prefer to use the median return rate (black line in Fig. 2D). This median return rate decreases from a value intermediate between the individual species’ return rates at short times *t* to asymptotic resilience at longer times *t*.

The example of Fig. 2 provides valuable intuitions about the recovery dynamics, also for systems with interacting species. However, one of these intuitions, namely that asymptotic resilience provides a lower limit for the recovery trajectories, does not hold generally. This is illustrated in Fig. 3, where we analyze another two-species system. The community matrix 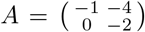 displays strongly asymmetric species interactions. All trajectories eventually return to equilibrium with rate R_∞_ = 1 (Fig. 3B). The short-term return to equilibrium has a much richer behavior. Many trajectories have short-term return rates either well above asymptotic resilience, or much smaller and even negative return rates (i.e., they are moving away from equilibrium). The latter phenomenon occurs because the system is reactive (Neubert & Caswell, 1997; systems with strongly asymmetric interactions are often reactive). Reactivity guarantees that there exist trajectories for which *R*_0_ < 0. However, it does not exclude that other trajectories have positive initial return rate *R*_0_. For the system in Fig. 3, for example, the distribution of R_0_ is mainly concentrated on positive values (Fig. 3D).

**Figure 3:**
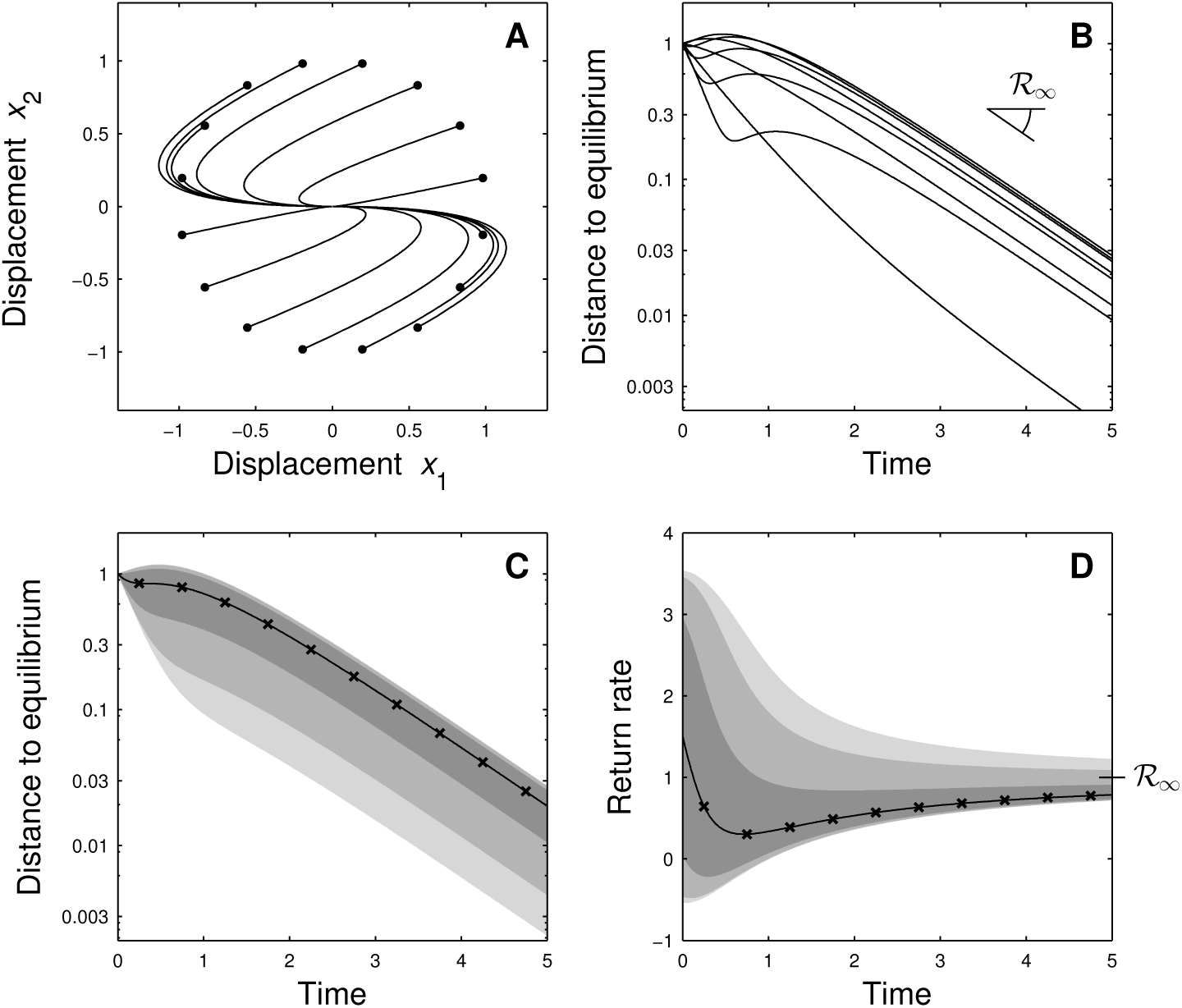
Return to equilibrium depends on perturbation direction – reactive case. Two-species system with community matrix 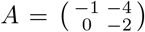. Panel A: Phase-plane trajectories for several perturbations *u*. Panel B: Dynamics of distance to equilibrium. For some perturbations the system initially moves away from the equilibrium, but for all perturbations the distance to equilibrium eventually decays at a rate equal to asymptotic resilience 𝓡_∞_. Panel C and D: Statistics of distance to equilibrium and of return rate 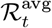 for random perturbations (fixed intensity, uniformly distributed). Full line: median computed from simulations; ×-marks: analytical approximation for median; shades of gray: 5%, 10%, 25%, 75%, 90% and 95% percentiles.

Generally, the distribution of return rates 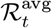 over time has a funnel shape: a broad distribution for small times *t* and an increasingly narrow distribution for larger times. This can be understood by looking at the initial and asymptotic return rates *𝓡*_0_ and *𝓡* _∞_. The distribution of 𝓡_0_ depends on the set of eigenvalues of the community matrix and on its reactivity (more precisely, on the set of eigenvalues of the symmetric part of the community matrix). Because these eigenvalues can span a large range, the distribution of 𝓡_0_ is typically wide. In contrast, the distribution of 𝓡_∞_ is extremely peaked at a single value, equal to asymptotic resilience. The distribution of return rates 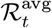 for 0 < *t* < ∞ connects these two extremes, giving rise to the characteristic funnel shape. In Appendix D we show that other stability measures based on return rates exhibit similar patterns.

## Averaging over perturbation directions

An ecosystem's return to equilibrium is not only determined by its dynamics, but also depends on the perturbation that initiated the displacement from equilibrium. This dependence has to be taken into account when predicting ecosystem recovery. However, often we do not know how exactly a perturbation, whether natural or experimentally induced, displaces the ecosystem. Here we propose a minimalistic way to deal with this uncertainty, and study the corresponding predictions for the recovery dynamics.

To model the uncertainty of the perturbation direction, we take a stochastic approach. We model the perturbation direction as a random variable, so that the return trajectories are also randomly distributed. Each realization of this randomness corresponds to a particular perturbation, which initiates a single return trajectory. Then, to obtain a practically relevant prediction, we propose to average the system response over the perturbation directions. Concretely, we construct a ‘typical’ return trajectory by taking, at each time after the perturbation, the average over the perturbation directions. Note that this typical trajectory is not necessarily the response to a particular perturbation. Rather, it is the composition of the average displacements through time.

In Appendix D we derive simple and accurate formulas for the median system response, which is easier to approximate than the average. These formulas depend on the system dynamics through the community matrix A and on the randomness of perturbation u encoded in a covariance matrix *C*. Component *C_ii_* of this matrix is the variance of initial displacement *u_i_* of species *i*. Component *C*_ij_ is the covariance of *u_i_* and *uj*; this covariance accounts for the fact that species *i* and *j* may undergo similar initial displacements. The approximate expressions for the median distance to equilibrium and the median return rate are then given by

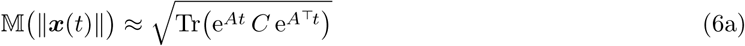

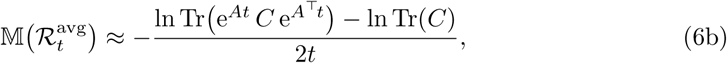

where the symbol 𝕄 stands for the median over the ensemble of perturbation directions. These expressions are easy to evaluate numerically (recall that Tr(A) denotes the trace, *e^A^* the matrix exponential and A^⊤^ the transpose of matrix A). In Appendix D we give analogous expressions for other stability measures based on return rates.

To illustrate their accuracy, we apply equations (6) to a few examples. First, we reconsider the examples of Figs. 2 and 3. We assume that the perturbation directions are uniformly distributed (on the unit circle, i.e., on the set of vectors *u* with ||*u*|| = 1; see Figs. 2A and 3A). This assumption corresponds to setting the perturbation covariance matrix *C* proportional to the identity matrix (*C_ii_* = 1/n and *C_ij_* = 0, with n the number of species in the system; see Appendix E). The agreement between the numerically computed medians (full line) and their analytical approximations (x-marks) is excellent (see Figs. 2D, 3C and 3D).

In the absence of additional information, the uniform distribution is an appropriate model for the perturbation randomness. As explained before, we only have to consider perturbation directions *u* with ||u|| = 1 (because perturbation intensity ||*u*|| does not affect return rates) and there is no reason to prefer one perturbation direction over another. However, there is additional information in the form of the equilibrium biomasses *N^*^_i_* (recall that the community matrix A is obtained by linearizing around the equilibrium vector N*). When species biomasses differ strongly, which is almost always the case in practice, the distribution over perturbation directions should be non-uniform.

To make this point clear, let us take a numerical example. Suppose a perturbation acts on a two-species system, in which the first species is ten times more abundant than the second species (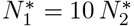). Compare then perturbation ‘a’ that mostly displaces species 1 (e.g., *u*_1_ = 10*u*_2_) and perturbation ‘b’ that mostly displaces species 2 (e.g., *u*_1_ *=* 0.1 *u*_2_; these perturbations are represented in Fig. 2). Perturbation ‘a’ affects both species equally in relative terms, while perturbation ‘b’ has a very strong effect on the rare species (in relative but also in absolute terms). Clearly, perturbation ‘a’ is more likely than perturbation ‘b’. This implies that the distribution over perturbations directions should assign a larger weight to perturbation ‘a’ than to perturbation ‘b’. This requirement disqualifies the uniform distribution as a suitable perturbation model.

There is no unique perturbation model in the case of an uneven abundance distribution. Here we propose to take the expected displacement *u_i_* of species *i* proportional to its equilibrium biomass *N^*^_i_*. That is, all species are perturbed equally in relative terms (i.e., relative to their equilibrium biomass). In Appendix E we prove that this assumption corresponds to setting the perturbation covariance matrix C proportional to the diagonal matrix of the squared species biomasses (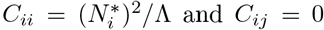, with 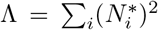). If all species have the same equilibrium biomass, we recover the formula for uniformly distributed perturbation directions. We use the biomass-dependent perturbation model in all the examples below.

In Fig. 4 we apply this model to the example of Fig. 2, assuming that species have different equilibrium biomass. The biomass of species 1, which recovers four times faster than species 2, is ten times larger than the biomass of species 2. Due to its larger biomass, species 1 is typically displaced more strongly than species 2. Hence, the perturbations are no longer uniformly distributed (as was the case previously, see Fig. 2A), but are concentrated close to the *x*_1_-axis corresponding to species 1 (see Fig. 4A). This implies that the fast recovery of species 1 has a much larger contribution to the average system recovery than in the previous scenario. For example, the median distance to equilibrium drops to about 5% of the initial displacement at the fast return rate of species 1 (Fig. 4C, for times *t <* 1). The slow return rate of species 2, equal to asymptotic resilience, governs the ecosystem response only later. As a result, one would probably only observe the fast recovery because the slow recovery corresponds to displacements that are too small to be detectable in practice. Note again the excellent agreement between numerical computations and analytical approximations (see Figs. 4C and 4D).

**Figure 4:**
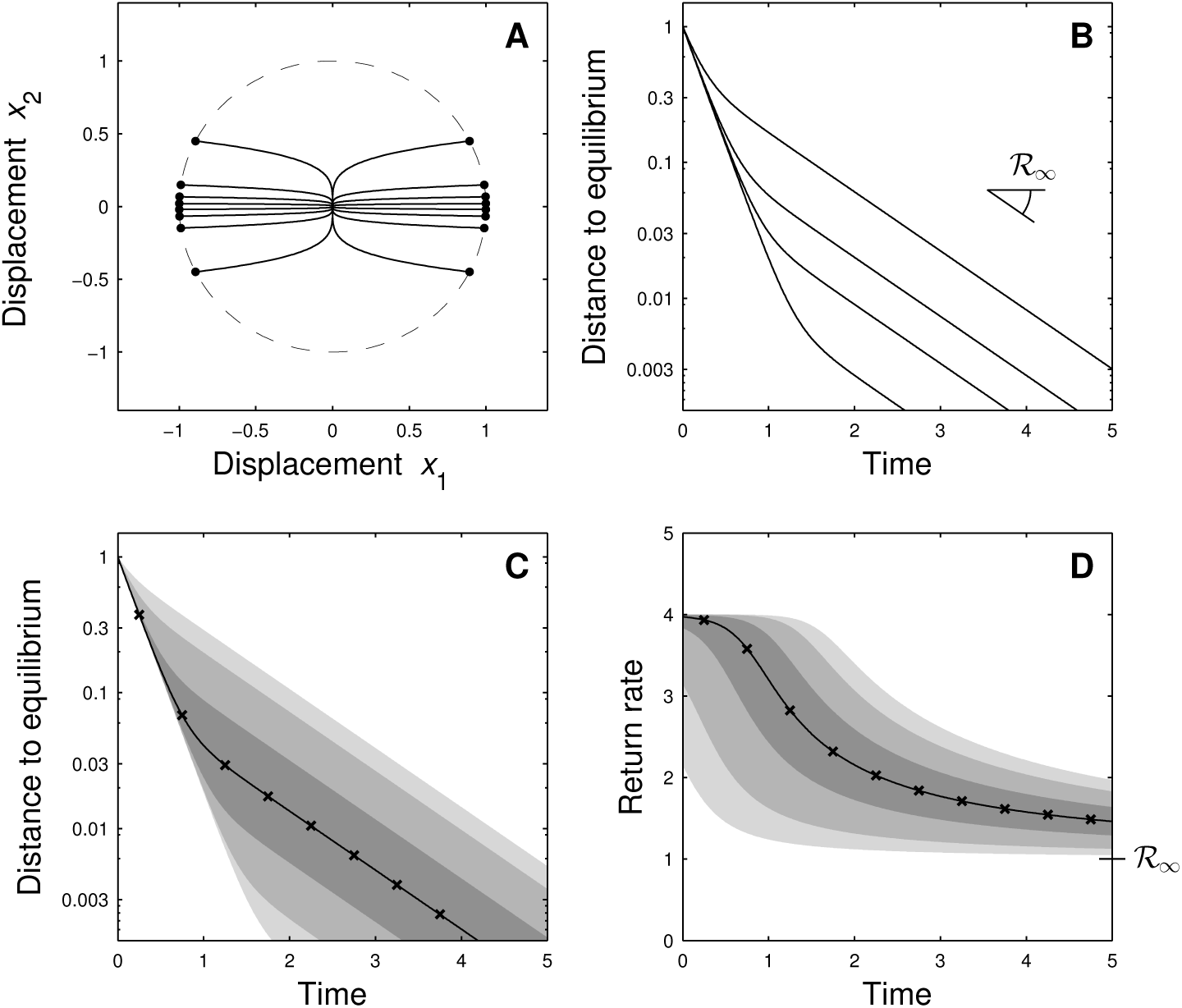
Return to equilibrium for biomass-dependent perturbations. Same system as Fig. 2, but here we take into account that perturbations affect abundant and rare species differently. Specifically, we assume that the equilibrium biomass of species 1 (the species with the fastest response) is ten times larger than the equilibrium biomass of species 2. As explained in the main text, this implies that a perturbation will typically displace mostly species 1. Perturbations are no longer spread out on the unit circle (panel A, dashed line), but tend to be directed along the *x*_1_-axis corresponding to species 1 (panel A, black dots). As a result, perturbations like the one labeled ‘a’ in Fig. 2 contribute more strongly to the statistics for random perturbation directions than perturbations like the one labeled ‘b’ in Fig. 2. For example, for almost all perturbations the distance to equilibrium becomes small (below 10% of the pulse perturbation) at a rate equal to the return rate of species 1 (rather than the return rate of species 2, which is equal to asymptotic resilience).

In summary, we have presented explicit predictions for the typical recovery dynamics that take into account uneven species abundance distributions. The latter information strongly affects the predicted return to equilibrium (compare Figs. 2 and 4). This framework can integrate additional information, such as a higher or lower vulnerability to perturbations of particular species, and positive or negative correlations in the responses of certain pairs of species.

## Effect of rare species on recovery dynamics

In the previous section we contrasted the immediate response of abundant and rare species to a pulse perturbation. We argued that on average rare species experience smaller biomass displacements, and hence contribute less to the short-term recovery dynamics. Moreover, we showed in an example (Fig. 4) that rare species can dominate the ecosystem response in the long term. This happens because rare species have the potential to introduce slow return rates in the system dynamics, and hence to determine asymptotic resilience. Here we explain why we expect this phenomenon to be common in real-world communities.

As a preliminary, we emphasize that there is no mathematically inevitable link between species rarity and long-term return rates. This can be easily illustrated using a system of non-interacting species. Suppose that species biomasses *N_i_* obey logistic growth,

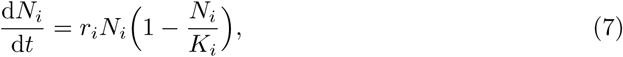
with intrinsic growth rate *r_i_* and carrying capacity *K_i_.* Due to the absence of interactions, each eigenvalue of the linearized community dynamics can be attributed to a different species. The eigenvalue associated with species *i* is equal to −*r_i_*. This shows that different parameters determine equilibrium biomass (*K*_i_) and eigenvalue (*r*_i_). Hence, by choosing parameter values appropriately, any species can provide the dominant eigenvalue, irrespective of abundance.

Consequently, the claim that rare species govern the long-term recovery cannot hold in full generality. However, we now argue that it can be expected as a common trend. To construct the argument, we restrict our attention to a particular type of rare species, namely those that play a minor role in the community. We call these species ‘satellite’, in opposition to ‘core’ species, which constitute the bulk of the community biomass. This terminology is borrowed from Hanski (1982), who introduced it to describe the regional distribution of species, whereas we apply it to the local level. Removing satellite species does not impinge on the community functioning. Satellite species do not affect the core species, or only weakly, but can be strongly affected by them. In particular, competition with core species prevents them from reaching higher abundances. Natural communities almost always contain numerous rare species, and while some of them might be an essential part of the community, a large majority can be expected to be satellite.

Surprisingly, despite their minor role in the community, satellite species can be predominant in the long-term return dynamics, as illustrated in Fig. 5. For simplicity we describe the coupled dynamics of a satellite species and the core community using a two-species competitive system, by aggregating the core species in a single biomass variable (see Appendix F for details). When the satellite species is absent, the return rate is a constant (Fig. 5, black dashed line), which we denote λ_0_.

**Figure 5:**
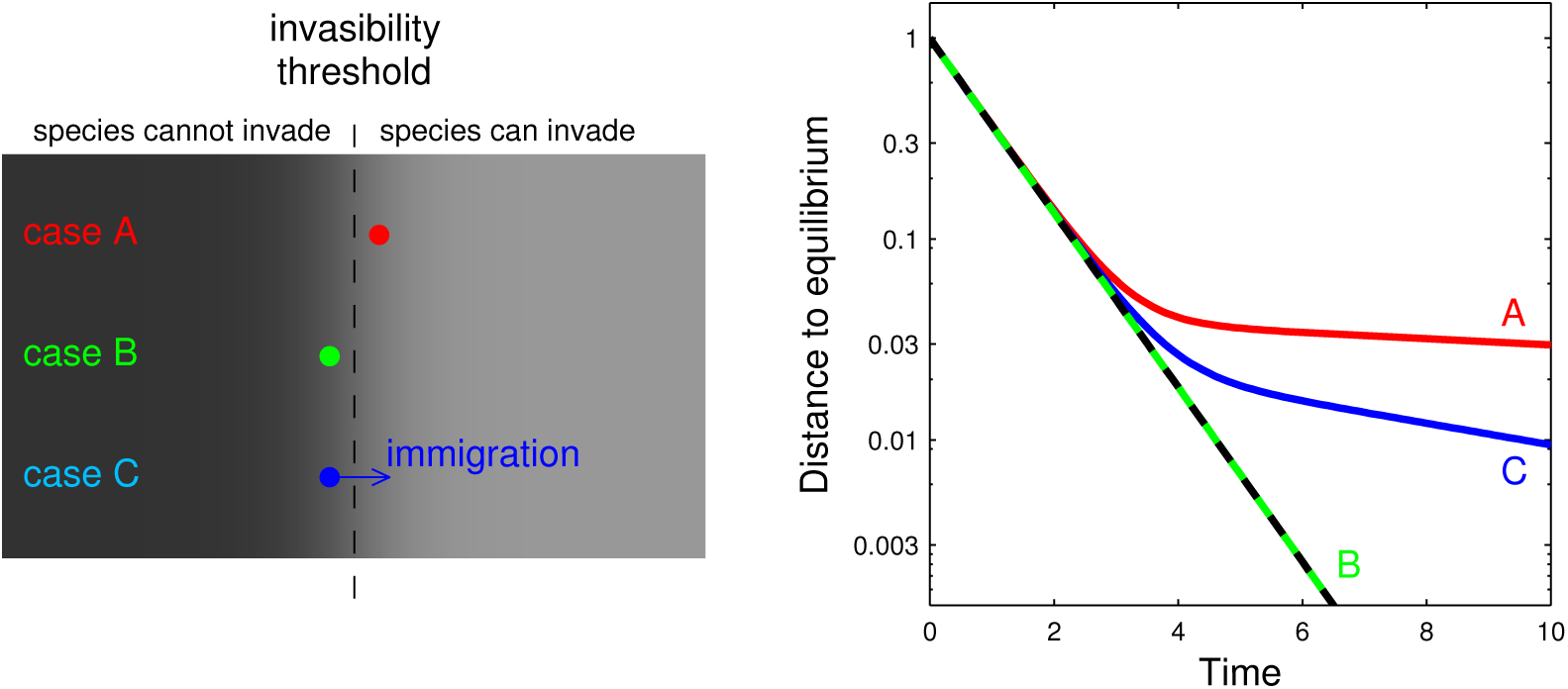
Effect of rare species on the long-term return to equilibrium. We study a two-species competitive Lotka-Volterra model, describing the introduction of a satellite species into an established community (see Appendix F). The system without that species has constant return rate (black dashed line in right-hand panel). Case A: If the introduced species has invasion fitness just above the invasibility threshold, it persists at a small equilibrium biomass. Compared to the community before the introduction, the short-term return to equilibrium does not change, but the long-term return to equilibrium becomes much slower (red line in right-hand panel). Case B: If the introduced species has invasion fitness just below the invasibility threshold, it cannot persist. As we consider perturbations that do not affect species that are absent at equilibrium, the return to equilibrium is the same as for the community before the introduction (green dashed line in right-hand panel). Case C: Instead of a closed community as in case B, we assume weak immigration, maintaining the introduced species at a small equilibrium biomass (source-sink dynamics). As in case A, the long-term return to equilibrium is much slower (blue line in right-hand panel). Model details and parameter values are given in Appendix F.

The presence of the satellite species modifies the recovery dynamics (Fig. 5, red line). The short-term recovery is not affected, but once the distance to equilibrium has decayed to a small fraction (≈ 5%) of the initial displacement, the return to equilibrium becomes much slower, corresponding to the asymptotic resilience of the coupled system (equal to 0.04 λ_0_ in the example, see Appendix F). Note that, in practice, this second part of the return dynamics, corresponding to small displacements from equilibrium, would be difficult to observe.

In natural communities species are often maintained by immigration, especially rare ones. To show the robustness of the previous argument, suppose that the satellite species is maintained in the community by immigration (i.e., a sink population). As before, the presence of the satellite species does not affect the short-term recovery, but it drastically slows down the long-term recovery (Fig. 5, blue line). Again, the part governed by asymptotic resilience (now equal to 0.11 λ _0_) sets in at very small displacements, and is therefore of limited practical interest.

The figure illustrates how the presence of a rare species can slow down the long-term return to equilibrium. This observation can be explained by looking at the dominant eigenvalue of the linearized dynamics, before and after introducing the satellite species. Because the satellite species has a negligible effect on the core community, the dynamics of the latter are essentially unaffected, and the eigenvalues of the core community without the satellite species are still eigenvalues of the coupled system including the satellite species. The coupled system has one additional eigenvalue, associated with the dynamics of that species. This eigenvalue can introduce a slow return rate (i.e., have small negative real part), especially if the satellite species is close to the invasibility threshold (see Fig. 5 and Appendix F), and can yield the dominant eigenvalue of the coupled system dynamics. When asymptotic resilience is determined by a single rare species, it contains limited information about community stability.

Each satellite species can provide the dominant eigenvalue, and we expect that real-world communities contain many such species. Hence, the influence of rare species on the long-term recovery dynamics might be widespread. We provide support for this claim using a random many-species community model. We impose that the equilibrium community has a realistic abundance distribution, with numerous rare species. The dynamics of species biomasses *N_i_* are governed by the competitive Lotka-Volterra equations,

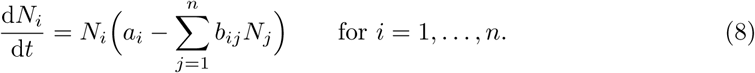

Parameter values of the *n* = 10 species are chosen as follows. First, we randomly generate the species biomasses *N^*^_i_* using a broken-stick model (MacArthur, 1957; Sugihara, 1980). We divide the total biomass 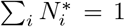 over the species by first allocating a random fraction (uniformly in the interval [0, 1]) of the total biomass to the first species, then by allocating a random fraction (uniformly in the interval [0, 1]) of the remaining biomass to the second species, and so on. Second, we randomly draw the competition coeffcients *b_ij_*: the intraspecific competition coefficients b_ii_ from the uniform distribution on the interval [0.5, 1], and the interspecific competition coefficients *b_ij_* with *i* ≠ *j* from the uniform distribution on the interval [0, 0.5]. Third, we determine the intrinsic growth rates *a_i_* such that the species biomasses *N^*^_i_* correspond to an equilibrium, that is, 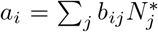. We check whether this equilibrium is stable, and discard the model realization if this is not case (which occurs for 23% of the model realizations).

The distribution of the recovery trajectories are shown in Fig. 6A. At time *t* = 100 most trajectories have decayed to a small fraction (≈ 5%) of the initial displacement. This level of displacement is typically no longer observable in noisy time series. However, the return rate continues to decrease, from 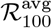 with median 0.02 to with median 0.0002 (Fig. 6B; note that the median 𝓡_∞_ corresponds to a horizontal line in Fig. 6A). By inspecting individual model realizations, we see that the disparity between 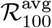 and 𝓡_∞_ is often associated with a rare species. In particular, when removing this species, the recovery dynamics up to time t = 100 do not change, while asymptotic resilience does (Fig. F.1). This is consistent with case A of Fig. 5. Hence, asymptotic resilience is determined by the specificities of rare species, which have almost no effect on the observable part of the recovery dynamics. This is further illustrated in Fig. 6C, where we show that, surprisingly, return rates 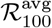 and 𝓡_∞_ have a weakly negative correlation. Although this negative correlation is due to the particular model randomness (and is not generally valid), it clearly illustrates that asymptotic resilience is an unreliable predictor for empirically relevant return rates.

**Figure 6.**
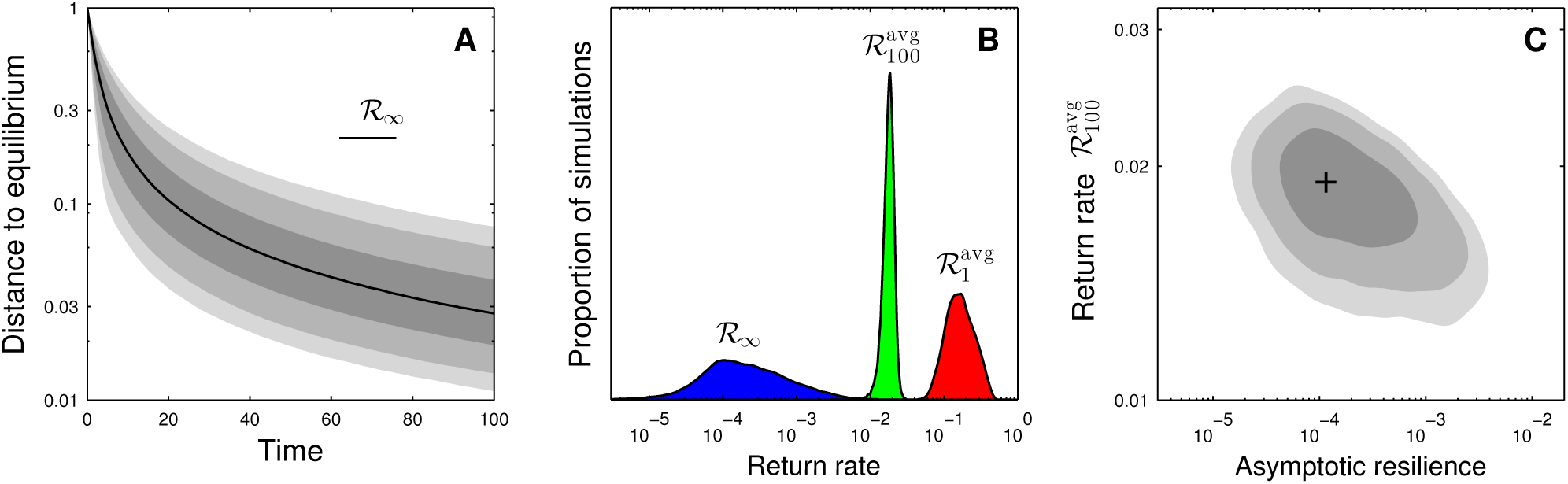
Return to equilibrium in a random community model. We analyze a Lotka-Volterra model with random competitive interactions. The equilibrium species biomass distribution is generated by the broken-stick model (see main text). Panel A: Statistics of distance to equilibrium for random model realizations (averaged over perturbation direction). Black line: median; shades of gray indicate 5%, 10%, 25%, 75%, 90% and 95% percentiles. Median asymptotic resilience 𝓡_∞_ corresponds to a virtually horizontal line (represented in the top-right part of the panel). Panel B: Probability distribution of return rates 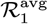, 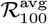 and 𝓡_∞_. Asymptotic resilience 𝓡_∞_ is orders of magnitude smaller than the finite-time return rates. Panel C: Joint probability distribution of return rates 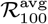 and 𝓡_∞_. Black cross: maximum; shades of gray indicate regions of 50%, 80% and 90% probability (corresponding to contour lines of the probability distribution). Asymptotic resilience 𝓡_∞_ is unreliable as a proxy for return rate 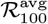. For this random community model there is even a (weakly) negative correlation between 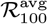 and 𝓡_∞_. The probability distributions in panels B and C were reconstructed using kernel density estimation on 10^4^ simulations.

In summary, while the short-term return dynamics are typically governed by the more abundant species, the return dynamics for longer times tend to be determined by rare species. This shift from abundant to rare species follows from two observations. First, a pulse perturbation is expected to initially generate the largest biomass changes in the abundant species, simply because they have larger biomass to start with. Second, towards the end of the recovery process, the latter becomes independent of the perturbation direction; it is then determined by the least stable species (in the sense of being closest to the invasibility threshold, see Fig. 5), which is often rare. The fact that distinct sets of species determine the short-term and long-term return rates explains why the latter are often unrelated.

## Discussion

In this paper we investigated means to quantify stability properties of ecosystems based on their recovery from a pulse perturbation. Introducing stability measures in the form of return rates, we unfolded the recovery process along two dimensions: time and species abundances. We first explained that short-term and long-term return rates can differ widely. In particular, short-term return rates heavily depend on perturbation directions, i.e., on the precise way the perturbation is distributed over species, while the long-term return rate is the same for almost any perturbation direction. This constitutes a qualitative difference in recovery along the time axis. We then argued that perturbations should be expected to mostly affect, in absolute terms, abundant species, but that the rare species tend to be the most unstable ones, i.e., the slowest to respond to perturbations. This is a distinction along the species abundances axis. Taking this reasoning further, we concluded that rare species typically control the asymptotic regime, while abundant species dominate the short-term response.

As a corollary, the asymptotic rate of return to equilibrium, or asymptotic resilience, should not be used as a proxy for the short-term recovery. Nevertheless, theoretical work on the return to equilibrium has focused almost exclusively on asymptotic resilience. For example, return time is often defined as the reciprocal of asymptotic resilience (a practice that dates back to Pimm & Lawton, 1977, 1978). But this theoretical construct need not be related to the actual return time, that is, the time it takes for the system to recover from a perturbation, which is mainly determined by the short-term response. Many ecologists seem to have built an intuition about the return to equilibrium based on very simple systems, such as single species, for which the return rate is constant over time. However, as illustrated by the examples in this paper, only slightly more complex systems exhibit much richer return dynamics, during which the return rate can change dramatically. We showed that in large, complex communities, due to the presence of species with very different abundances, asymptotic resilience need not even be a good predictor of return rates at longer times. Similarly, because asymptotic resilience does not depend on the perturbation direction, many ecologists seem to assume that the same holds for the entire recovery process. This intuition is erroneous because, as we have shown, the short-term return rates can, and often do, strongly depend on the perturbation direction.

Previous work has stressed that the asymptotic regime is often not representative of the short-time dynamics (Hastings, 2004, 2010). This issue has been particularly well studied in population ecology. It is generally recognized that depending on initial conditions the population dynamics can be governed by transient effects, which are missed out when analyzing the asymptotic regime alone (Caswell, 2001; Ezard *et al.,* 2010). Practical tools are available to systematically investigate the transient dynamics of population models, and to incorporate these transient effects into predictions of future population dynamics (Caswell, 2007; Stott *et al.,* 2011). Clearly, there are close parallels with the findings reported in this paper. It would be worthwhile to scrutinize whether theoretical insights and practical tools developed by population ecologists can enrich the study of ecosystem stability.

Because our work emphasizes the importance of the short-term recovery, it is closely related to the work of Neubert & Caswell (1997). They studied the instantaneous return rate immediately after a pulse perturbation, and showed that it can be negative (i.e., the distance to equilibrium initially increases) even if the asymptotic return rate is positive (i.e., even if the system is stable). They introduced the term ‘reactive’ to denote systems for which this phenomenon occurs, and argued that many real-world systems can be expected to be reactive. However, we have shown that the initial return rate displays a particularly strong dependence on the perturbation direction. Therefore, the existence of a perturbation with a negative initial return rate does not imply that the initial return rate is negative for all or even most perturbations. For instance, in Fig. 3, the vast majority of perturbations are met with positive initial return rates, despite the system being reactive. In fact, it can be shown that the median initial return rate is always positive and larger than asymptotic resilience, both for non-reactive and reactive systems (see Appendix D). This suggests that the system property of being reactive does not provide much information about the initial return rate for an actual pertubation. The theory of reactive systems deals with the initial return rate for the worst-case perturbation, i.e., the perturbation producing the most negative initial return rate, but does not tell us how the system typically responds to a perturbation. By studying this typical response, our paper can be interpreted as an extension of Neubert & Caswell’s.

This paper strives to develop theory for empirically relevant stability measures. The longterm return to equilibrum is of limited practical interest, because it corresponds to small displacements, which are often indistinguishable inevitable fluctuations at the equilibrium state. Also, especially in field studies, the asymptotic response to a first perturbation might be concealed by the occurrence of a second one. Therefore, available empirical data are often restricted to the short-term recovery, which is explicitly addressed by our theory. Shortterm responses depend on the perturbation direction, and we argued that the most relevant predictions are obtained by averaging over the perturbation distribution. We derived accurate formulas for the median return rate as a function of the time elapsed since the perturbation. These formulas can be evaluated as easily as asymptotic resilience, to which the median return rate converges in the limit of very long times. Thus, our work provides a theoretical framework to study the transient recovery following perturbations and to predict return times to equilibrium in community and ecosystem models.

This theoretical framework depends on two technical assumptions. First, we assumed that the reference state, i.e., the state in which the ecosystem settles at the end of the recovery process, is an equilibrium. Alternatively, and more realistically, we could consider a fluctuating reference state. If these fluctuations are small compared with the displacement induced by the pulse perturbation, then they do not affect the analysis of the short-term recovery. Second, we assumed that the recovery trajectories remain close to equilibrium. This allowed us to rely on the theory of linear dynamical systems, which are widely used by both theorists and empiricists to describe and interpret ecological dynamics (Gurney & Nisbet, 1998; Caswell, 2001). The non-linear part of ecosystem dynamics is expected to make the difference between short-term and long-term return rates even greater. Indeed, non-linearities can have a strong effect on the short-term response, but leave the long-term response essentially unchanged, because the latter corresponds to small displacements for which the linear approximation is accurate. Hence, non-linearities are an additional source of discrepancy between short-term and long-term responses, and therefore reinforce our argument.

The integration of theoretical and empirical approaches has been identified as one of the main challenges for research on ecological stability (Ives & Carpenter, 2007; Donohue *et al.,* 2016). We have tried to make the theory of how ecosystems recover from pulse perturbations more practically relevant by emphasizing short-term responses. Future work could address how to translate our findings into concrete recommandations (see Table 1 for some preliminary suggestions). While restricted to pulse perturbations, our paper might inspire analogous studies for other stability measures, such as the response to press perturbations and the temporal variability of ecosystems (see Arnoldi *et al.,* 2016 and Haegeman *et al.,* 2016 for first steps in this direction).

**Table 1.**
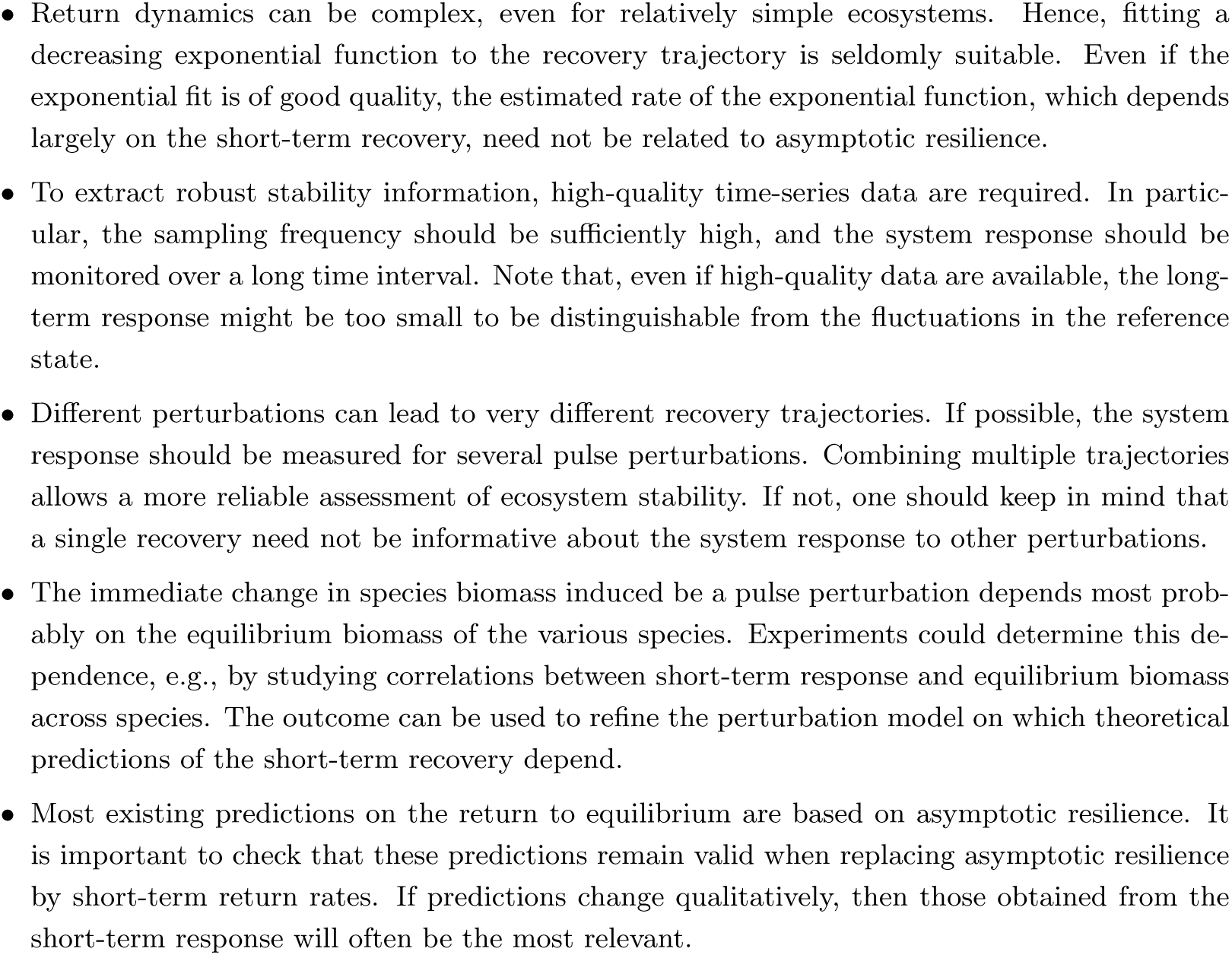
Suggestions to improve the connection between empirical and theoretical work on ecosystem recovery.

## Acknowledgements

We thank José Montoya for very helpful discussions, and Michael Cortez for constructive comments on the manuscript. This work was supported by the TULIP Laboratory of Excellence (ANR-10-LABX-41) and by the BIOSTASES Advanced Grant, funded by the European Research Council under the European Union’s Horizon 2020 research and innovation programme (grant agreement No 666971).

